# Regulation of Glypican 6-mediated Wnt activation maintains TDP-43 nuclear localization in neurons

**DOI:** 10.1101/2025.09.24.678385

**Authors:** Nan Zhang, Shanthini Sockanathan

## Abstract

Abnormalities in TDP-43 (Transactive response DNA-binding protein 43kDa) localization and function span multiple neurodegenerative diseases and are implicated in driving neuronal degeneration and loss. Nuclear pore complex (NPC) abnormalities and disrupted nucleocytoplasmic trafficking (NCT) contribute to TDP-43 mislocalization, but how these cellular changes are initiated in disease is unclear. Glycerophosphodiester phosphodiesterase 2 (GDE2) is a surface glycosylphosphatidylinositol (GPI)-anchor cleaving enzyme that encodes a physiological pathway that ensures NPC integrity, appropriate NCT, and nuclear TDP-43 expression and function in adult neurons by negatively regulating canonical Wnt signaling. Notably, studies of human postmortem tissue and patient-derived neuronal models suggest that the failure of GDE2-dependent regulation of Wnt contributes to TDP-43 abnormalities in disease. Here we show that GDE2 inhibits persistent neuronal Wnt activation by regulating the surface expression of the GPI-anchored protein, Glypican-(GPC)6. Excessive GPC6 surface expression potentiates neuronal Wnt activation *in vivo*, resulting in NPC disruption, alterations in Ran-dependent NCT, and TDP-43 mislocalization, while genetic reduction of GPC6 in mice lacking GDE2 rescues these cellular abnormalities. Thus, GDE2, GPC6, and the canonical Wnt pathway form a physiologically important signaling axis important for NPC integrity, appropriate NCT, and TDP-43 nuclear function in neurons that, when disrupted, may underlie associated neuropathologies in disease.

## Introduction

TDP-43 (Transactive response DNA binding protein 43 kDa) is a ubiquitously expressed protein that is required for the appropriate regulation, processing, and stability of thousands of transcripts^1–3^. In adult neurons, loss of TDP-43 nuclear function leads to neurodegeneration, suggesting that TDP-43 is critically required for the appropriate splicing and expression of genes essential for neuronal function and health^1,4^. Importantly, TDP-43 abnormalities such as TDP-43 cytoplasmic mislocalization, nuclear exclusion, aggregation, and downregulation are major hallmarks of various neurodegenerative diseases, including Alzheimer’s disease (AD) and AD-related dementias such as Amyotrophic Lateral Sclerosis (ALS) and ALS/Frontotemporal Dementia (FTD)^3,5–7^. These pathologies are accompanied by the loss of TDP-43 molecular function, which causes the incorporation of cryptic exons in TDP-43 target mRNAs, resulting in gene dysregulation and the synthesis of non-functional proteins^6,8–11^. Thus, TDP-43 dysfunction is thought to be a driver of neurodegeneration and loss across multiple neurodegenerative diseases. Several pathways are known to contribute to TDP-43 abnormalities in disease, including the erosion of the nuclear pore complex (NPC) and disrupted nucleocytoplasmic transport (NCT)^7,12,13^. However, the mechanisms by which these cellular changes in disease are initiated are still not well understood.

Glycerophosphodiester phosphodiesterase 2 (GDE2) is one of three six-transmembrane proteins that cleave the glycosylphosphatidylinositol (GPI) anchor that tethers some proteins to the outer leaflet of the plasma membrane^14–16^. GDE2 is expressed in neurons, a subset of terminally differentiated oligodendrocytes, and vascular endothelial cells, while GDE3 is expressed in oligodendrocyte precursor cells and astrocytes^17–21^. GDE6 is expressed in neural progenitors but has not been detected in mammals^22,23^. In the developing nervous system, neuronally expressed GDE2 regulates the differentiation of subsets of late-born spinal and cortical neurons and contributes to oligodendrocyte differentiation via the release of maturation signals^18,24,25^. GDE2 expression persists in the adult nervous system, where it has separate roles in regulating pathways that are important for neuronal function and survival^26–29^. Prominent among these roles is that GDE2 is essential for maintaining NPC integrity, appropriate NCT, and ensuring TDP-43 nuclear localization and function^29^. Mechanistic studies in mouse models and neurons derived from human induced pluripotent stem cells (iPSCs) determined that GDE2 is required to prevent the sustained activation of canonical Wnt signaling in neurons, which is sufficient to erode the NPC, impair NCT, and disrupt TDP-43 nuclear expression^29^. Thus, GDE2 is a physiological regulator of neuronal canonical Wnt activation that is important for preserving NPC, NCT, and TDP-43 nuclear expression and function. Further evidence suggests that the GDE2/Wnt regulatory pathway is relevant to TDP43 pathologies in disease. GDE2 aberrantly accumulates in intracellular compartments in the neurons of postmortem brains of patients with AD, ALS, and ALS/FTD^27,28^; strikingly, neurons with GDE2 accumulations correlate precisely with TDP-43 pathologies^29^. Further, iPSC-derived neurons from patients with ALS show a reduction in GDE2 protein and increased canonical Wnt activation, which, when dampened by pharmacological agents, partly rescues TDP-43 nuclear function^29^. Deeper insight into the mechanisms by which GDE2 regulates canonical Wnt signaling in mature neurons could help clarify initiating mechanisms that underlie NPC, NCT, and TDP-43 abnormalities in disease.

GDE2 acts at the cell surface to cleave the GPI-anchor of its substrates and release them from the plasma membrane^14–16^. Accordingly, we hypothesize that GDE2 regulates the cell surface expression and activity of GPI-anchored proteins that activate or potentiate Wnt signal transduction to regulate NPC integrity, NCT, and TDP-43 localization. Glypican (GPC)2, GPC4, and GPC6 are established substrates of GDE2^16^. They belong to the Glypican family of GPI-anchored heparan sulfate proteoglycans that undergo extensive post-translational modifications, including the attachment of glycosaminoglycan (GAG) chains and endoproteolytic cleavage by furin-like convertases^30^. Glypicans are evolutionarily conserved from *Drosophila* to humans. In mice and humans, there are six Glypican family members that are divided into two subfamilies: GPC1/2/4/6, which correspond to *Drosophila* Dally-like protein (Dlp), and GPC3/5, which are homologues of *Drosophila* Dally^30^. Between the two subfamilies, GDE2 can cleave GPC1/2/4/6 but not GPC3/5^16^. Dlp has been shown to potentiate Wnt activation at suboptimal concentrations and can bind to the lipid moieties of Wnt ligands, facilitate their transport across cells, and accordingly regulate their accessibility and binding to Wnt receptors^30–32^. Thus, Glypicans related to Dlp are important regulators of canonical Wnt activation. Interestingly, GPC1, GPC4, and GPC6 show impaired release in the spinal cord of the *SOD^G93A^* mouse model of familial ALS, which exhibits impaired GDE2 activity^26^. Moreover, mutations in *GPC6* are associated with AD and ALS in genome-wide association studies (GWAS), and GPC6 exhibits increased expression and altered localization in spinal cord tissues from patients with ALS with TDP-43 proteinopathies^33–37^. These observations suggest that the Dlp-class Glypicans are good candidates for substrates targeted by GDE2 to regulate Wnt-mediated effects on NPC, NCT, and TDP-43 localization in health and disease.

Here, we sought to determine the physiologically relevant Glypicans that are regulated by GDE2 to modulate canonical Wnt signaling in adult cortical neurons. We show that GDE2 regulates GPC6 surface expression in cortical neurons to control Wnt activation and that increased levels of GPC6 are sufficient to promote sustained canonical Wnt signaling, causing abnormalities in the NPC, NCT, and disrupting the nuclear localization of TDP-43. Thus, the GDE2/GPC6 regulation of Wnt activation is a physiological pathway that, when impaired, may contribute to NPC, NCT, and TDP-43 pathologies in disease.

## Results

### Gde2 and Gpc6 expression and regulation in cortical neurons

To identify candidate Glypicans that overlap with GDE2 expression in excitatory cortical neurons, we took advantage of publicly available single-cell RNA-sequencing (scRNAseq) datasets of the adult mouse cortex^38^, focusing on excitatory neurons that show *Gde2* expression. Consistent with previous studies, *Gde2* is expressed throughout the adult cortex, and is enriched in excitatory neurons in deep layers of the cortex corresponding to Layers 5 and 6^21^ (Fig. 1a). Of the 6 Glypican family members, *Gpc6* shows the highest mRNA expression in *Gde2*-expressing cortical excitatory neurons (Fig. 1a). *Gpc6* is expressed in subsets of neurons across all cortical layers, with robust expression in Layer 5 and Layer 6 excitatory pyramidal tract (PT) and intratelencephalic (IT) neurons. *Gpc1, 4,* and *5* are expressed at lower levels than *Gpc6* in subsets of deep-layer neuronal layers, with *Gpc*5 also showing expression in Layers 2/3. In contrast, *Gpc2* and *Gpc3* are not co-expressed with *Gde2* in cortical neurons.

**Figure 1:**
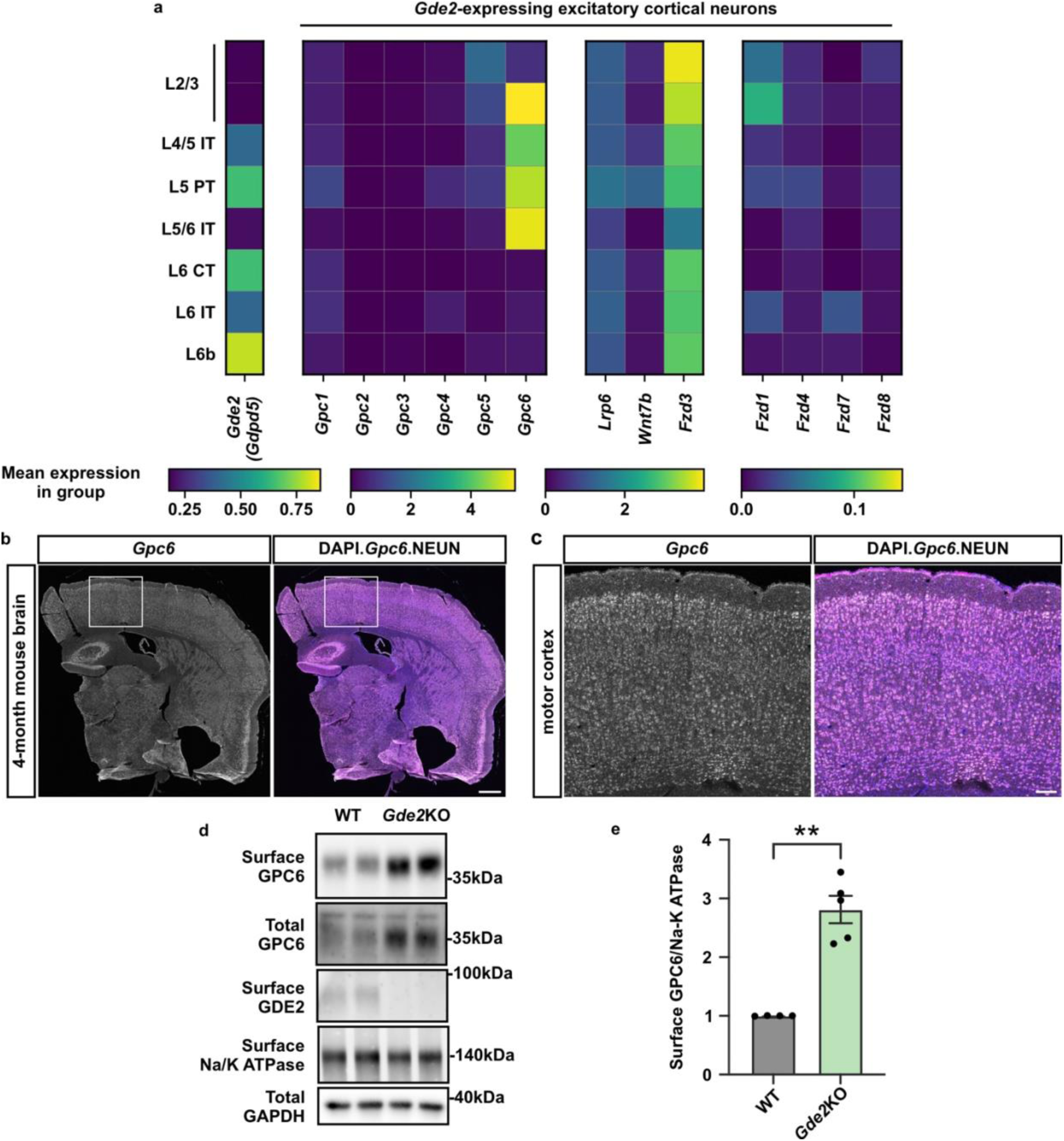
*Gpc6* is co-expressed with *Gde2* in the mouse cortex and regulated by *Gde2* expression. **a**. Heat map showing the expression of *Gde2* (*Gdpd5*) in excitatory neurons in the mouse cortex (left) and the expression of Glypican family proteins and Wnt signaling components in *Gde2*-expressing cortical neurons (right). Each subpanel is scaled differently to highlight the range of expression levels. Data from the Allen Brain Atlas. L: layer. **b-c**. Representative images of RNAscope probing for *Gpc6* transcripts in coronal sections of the adult mouse brain. The boxed area of the motor cortex in **b** is shown in magnified view in **c**. Scale bar: **b** = 500µm; **c** = 100µm. **d-e**. Representative western blot images of surface biotinylation of DIV21 WT and *Gde2*KO primary cortical neurons (**d**) and graph quantifying the surface levels of furin-converted GPC6 normalized to the membrane protein, Na-K ATPase (**e**). WT vs. *Gde2*KO: **p = 0.0015. n = 4 WT, 5 *Gde2*KO. Welch’s t-test. All graphs: mean ± sem.

We focused our analysis on GPC6 because of the level and extent of its co-expression with *Gde2* in cortical neurons. RNAscope experiments on sections of 4-month-old WT mice confirm that *Gpc6* is expressed throughout the brain and is distributed in neurons across all cortical layers (Fig. 1b-c). These observations, combined with the knowledge that GPC6 is an established physiological substrate of GDE2, suggest GPC6 as a potential candidate for mediating GDE2-dependent regulation of canonical Wnt signaling in excitatory neurons. GDE2 acts at the cell surface to cleave the GPI-anchor of its substrates and release them into the extracellular space. Accordingly, if GPC6 is a substrate of GDE2 in excitatory cortical neurons, loss of GDE2 should result in increased GPC6 surface expression and, conceivably, an increase in total levels of GPC6 protein. To test this idea, we performed surface biotinylation to examine the expression of GPC6 and GDE2 in cortical neurons prepared from WT and *Gde2*KO postnatal day (P) 0 pups that were cultured for 21 days in vitro (DIV). Western blot analysis showed that there is an increase in surface GPC6 expression and total amounts of GPC6 protein in *Gde2*KO neurons compared with WT (Fig.1 d-e). Taken together, these observations are in line with the model that GDE2 regulates GPC6 surface expression to suppress sustained canonical Wnt signaling in excitatory cortical neurons in the adult brain.

### GPC6 potentiates Wnt7b mediated signaling

Wnt ligands comprise a large family of proteins, and to date, 19 Wnt ligands have been identified in mice^39–41^. We analyzed the expression of Wnt ligands in the same scRNAseq data described above and found that *Wnt4, Wnt9a,* and *Wnt7b* are co-expressed with *Gde2* and *Gpc6* in excitatory cortical neurons (Fig. 1a and Fig. S1a). Wnt4 has been reported as a non-canonical Wnt ligand^42^, while Wnt9b is implicated in both canonical and non-canonical Wnt signaling pathways^43^. Wnt7b typically mediates canonical Wnt signaling^44,45^. Because GDE2 is known to regulate canonical Wnt signaling in cortical neurons^18,29^, we examined the relationship between GPC6 and Wnt7b-dependent canonical Wnt signaling. To examine if GPC6 is capable of potentiating Wnt7b-mediated signaling, we utilized SH-SY5Y cells, a neural-like immortalized cell line responsive to Wnt activation. To facilitate the detection of GPC6, we generated an N-terminal HA-tagged GPC6 (HA-GPC6). Notably, addition of the HA tag did not interfere with GPC6 processing by furin convertases (Fig. 2a), GPC6 trafficking to the cell surface (Fig. S2a-b), or GPC6 release by GDE2^46^. We transfected SH-SY5Y cells with a low amount of plasmid expressing Wnt7b or GFP to establish conditions where Wnt7b expression did not alter basal levels of non-phosphorylated (p)-β-catenin amounts, consistent with suboptimal Wnt pathway activation (Fig. 2a-b, grey bars). Under the same conditions, co-transfection of Wnt7b with GPC6 led to a significant increase in the amounts of non-p-β-catenin; however, no synergy was observed when GPC6 was co-transfected with a control plasmid expressing GFP (Fig. 2a-b, green bars). These observations support the notion that increased surface expression of GPC6 is sufficient to potentiate canonical Wnt7b-mediated signaling. Interestingly, total levels of GPC6 were reduced in the presence of Wnt7b (Fig. 2a), consistent with previous studies that glypicans are internalized with the Wnt-receptor complex^47,48^, which we speculate is targeted for degradation.

**Figure 2.**
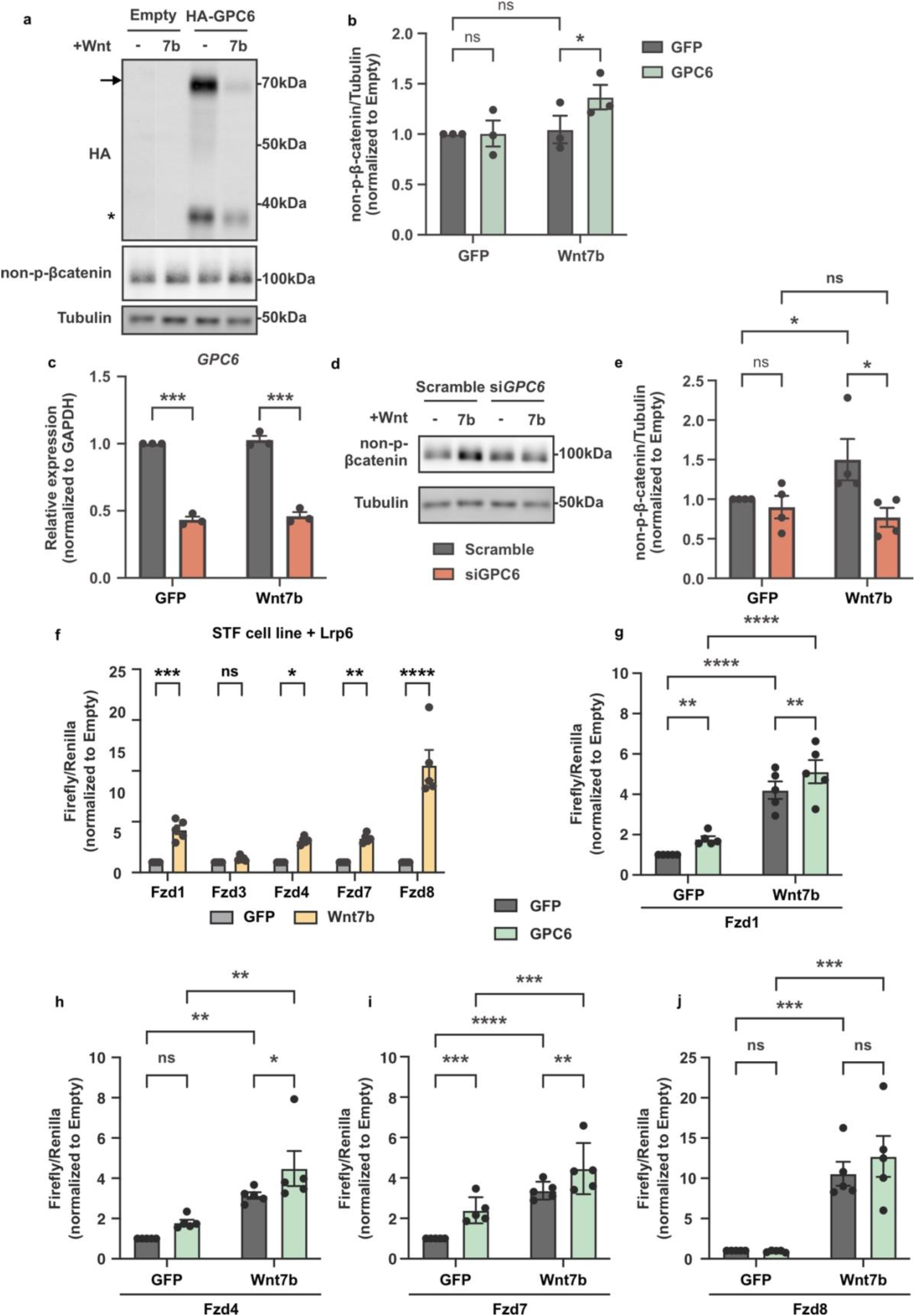
GPC6 potentiates Wnt7b signaling transduction. **a-b.** Representative western blots (**a**) of SH-SY5Y cells transfected with plasmids expressing HA-GPC6 and Wnt7b and graph (**b**) quantifying non-p-β-catenin levels normalized to Tubulin. Arrow marks full-length HA-GPC6; Asterisk marks N-terminal HA-GPC6 protein after cleavage by furin convertase. GFP-GFP vs. GFP-GPC6: ns = 0.9449; Wnt7b-GFP vs. Wnt7b-GPC6: *p = 0.0253; GFP-GFP vs. Wnt7b-GFP: ns = 0.7818. RM two-way ANOVA. n = 3, mean ± s.e.m. **c**. Graph quantifying *GPC6* mRNA levels treated with siRNAs. GFP-scramble vs. GFP-si*GPC6*: ***p = 0.0003; Wnt7b-scramble vs. Wnt7b-si*GPC6*: ***p = 0.0003. RM two-way ANOVA with Sidak’s multiple comparison test. n = 4, mean ± s.e.m. **d**. Representative western blots of SH-SY5Y cells transfected with plasmids expressing Wnt7b and treated with siRNAs. **e**. Graph quantifying non-p-β-catenin protein levels normalized to Tubulin. GFP-scramble vs. GFP-si*GPC6*: ns = 0.8438; GFP-scramble vs. Wnt7b-scramble: *p = 0.0372; Wnt7b-scramble vs. Wnt7b-siGPC6: *p = 0.0338; GFP-siGPC6 vs. Wnt7b-siGPC6: ns = 6939. RM two-way ANOVA. n = 4, mean ± s.e.m. **f-j**. Topflash luciferase assay in STF cells quantifying Firefly luciferase activity, reporting Wnt transcriptional activity, normalized to Renilla expressed under a universal promoter. **f.** GFP vs. Wnt7b with Fzd1: ***p = 0.0003; Fzd3: ns p = 0.9898; Fzd4: *p = 0.0207; Fzd7: **p = 0.0086; Fzd8: ***p < 0.0001. Ordinary two-way ANOVA with Sidak’s multiple comparison test, with a single pooled variance. n = 5, mean ± s.e.m. **g:** Fzd1: GFP-GFP vs. GFP-GPC6 **p = 0.0034; GFP-GFP vs. GFP-Wnt7b ****p < 0.0001; Wnt7b-GFP vs. Wnt7b-GPC6 **p = 0.0019; GFP-GPC6 vs. Wnt7b-GPC6: ****p < 0.0001. **h:** Fzd4: GFP-GFP vs. GFP-GPC6 ns p = 0.1370; GFP-GFP vs. GFP-Wnt7b **p = 0.0076; Wnt7b-GFP vs. Wnt7b-GPC6 *p = 0.0358; GFP-GPC6 vs. Wnt7b-GPC6: **p = 0.0034. **i:** Fzd7: GFP-GFP vs. GFP-GPC6 ***p = 0.0008; GFP-GFP vs. GFP-Wnt7b ****p < 0.0001; Wnt7b-GFP vs. Wnt7b-GPC6 **p = 0.0020; GFP-GPC6 vs. Wnt7b-GPC6: ***p = 0.0002. **j:** Fzd8: GFP-GFP vs. GFP-GPC6 ns p = 0.9327; GFP-GFP vs. GFP-Wnt7b ***p = 0.0004; Wnt7b-GFP vs. Wnt7b-GPC6 ns p = 0.0644; GFP-GPC6 vs. Wnt7b-GPC6: ***p = 0.0002. RM-two-way ANOVA. n = 5, mean ± s.e.m.

To test if GPC6 is required to promote Wnt7b signaling, we took advantage of the observation that SH-SY5Y cells express endogenous GPC6^46^, which likely contributes to basal levels of Wnt activation determined by non-p-β-catenin expression (Fig. 2a). Treatment of SH-SY5Y cells transfected with plasmids expressing GFP or Wnt7b with siRNAs targeting *GPC6,* led to an approximately 50% reduction of endogenous *GPC6* mRNA expression when compared to control siRNAs (Fig. 2c). Notably, GPC6 knockdown ablated the Wnt7b-dependent increase in non-p-β-catenin that was observed in control siRNA treated conditions (Fig. 2d-e), indicating that endogenous GPC6 is required for Wnt7b-dependent canonical Wnt activation. Taken together, these observations suggest that GPC6 is necessary and sufficient to facilitate Wnt7b signal transduction.

We next examined the surface components that mediate Wnt7b signaling in concert with GPC6. Notably, β-catenin also plays major roles at adherens junctions and links cadherin cell adhesion molecules to the actin cytoskeleton^49^. To focus our analysis on nuclear β-catenin relevant to Wnt7b-dependent activation of transcription, we utilized a heterologous system using the Wnt reporter cell line, SuperTopflash (STF)^44^. STF is a stable HEK293T cell line expressing a firefly luciferase reporter gene under the regulation of 7 TCF/LEF binding sites, which is activated when non-p-β-catenin binds to TCF/LEF transcription factors in response to canonical Wnt activation^39,44,50^. Analysis of the sc-RNAseq data described in Figure 1 showed that the Wnt receptors *Fzd1*, *Fzd3*, *Fzd4*, *Fzd7*, and *Fzd8,* and the Wnt co-receptor *Lrp6* are co-expressed with *Gde2* and *Gpc6* in deep-layer cortical neurons (Fig. 1a and Fig. S1b). STF cells do not express Lrp6; accordingly, we co-expressed Wnt7b, Lrp6, and each identified Fzd receptor in STF cells and assayed cells for luciferase activity (Fig. 2f). We found that Wnt7b activates canonical Wnt signaling when co-transfected with Lrp6 and Fzd1, Fzd4, Fzd7, or Fzd8 (Fig. 2f), but not Fzd3. This is consistent with previous reports that Fzd3 mainly mediates non-canonical Wnt signaling^44^. Next, we tested whether GPC6 potentiates Wnt7b activation when co-expressed with Fzd1, Fzd4, Fzd7, and Fzd8 in this paradigm. We found that GPC6 is capable of potentiating Wnt7b activation when co-expressed with Fzd1, Fzd4, and Fzd7 but not Fzd8 (Fig. 2g-j). Moreover, we observed a modest increase in Wnt activation when GPC6 is overexpressed with Fzd1 and Fzd7 in the absence of exogenous Wnt7b expression, suggesting that GPC6 can potentiate endogenous Wnt signaling in STF cells (Fig. 2g and i). Taken together, these *in vitro* observations suggest that GPC6 can potentiate canonical Wnt7b/Lrp6 activation via a subset of Fzd receptors that include Fzd1, Fzd4, and Fzd7, all of which are co-expressed with GDE2 in deep-layer cortical neurons.

### GPC6 overexpression in the cortex phenocopies *Gde2*KO

Previous studies show that GDE2 prevents the sustained activation of canonical Wnt signaling in neurons to ensure NPC integrity, appropriate NCT, and TDP-43 nuclear expression^29^. We hypothesize that increased surface GPC6 expression in neurons elicited by GDE2 loss causes aberrant Wnt activation that drives NPC, NCT, and TDP-43 abnormalities. To test this hypothesis, we utilized a transgenic Wnt reporter mouse line that contains a H2B-MYC-GFP fusion reporter under the control of TCF/LEF1 binding sites integrated into the *Rosa26* genetic locus (WT;*Wnt-MYC-GFP*)^45^, to examine whether GPC6 overexpression can activate neuronal Wnt signaling *in vivo*. In these animals, Wnt activation can be visualized at single-cell resolution by MYC or GFP expression. We generated adeno-associated virus (AAV) expressing HA-GPC6 (AAV HA-GPC6) or control virus (AAV HA-GFP-MYC or AAV HA-GFP). We injected WT;*Wnt-MYC-GFP* mice at P28 with AAV HA-GPC6, AAV HA-GFP, or AAV-HA-GFP-MYC by the retro-orbital route and examined animals 10 days later for Wnt activation by immunostaining cortical neurons for MYC expression. We found that cortical neurons expressing HA-GPC6 showed an increase in nuclear MYC expression compared to neurons expressing HA-GFP, suggesting that GPC6 overexpression induces canonical Wnt signaling in neurons *in vivo* (Fig. 3a-b).

**Figure 3:**
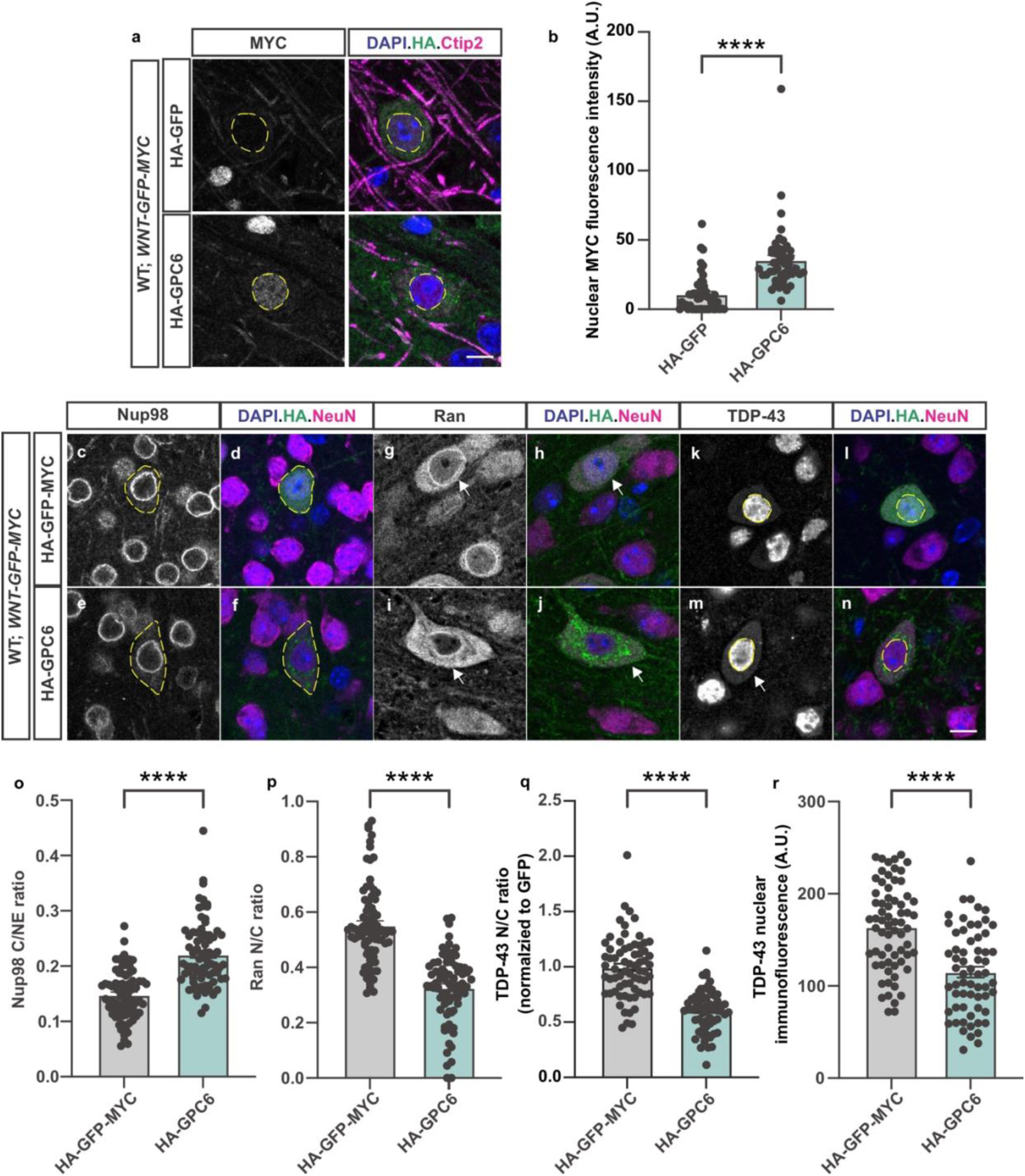
GPC6 overexpression activates Wnt signaling and drives NPC, NCT, and TDP-43 abnormalities. **a, c-n**: Representative images of immunostained cortical sections of AAV-transduced WT;*Wnt-GFP-MYC* animals 10 days post-injection. Dashed lines highlight the nuclei of cells transduced with AAV HA-GFP or AAV HA-GPC6 in **a** and **k-n**; dashed lines outline the cell body of transduced cells in **c-f**; arrows highlight transduced cells in **g-j**. Scale bar: **a**, **c-n**: 10μm. **b**: Graph quantifying nuclear MYC immunofluorescence intensity in neurons expressing HA-GFP or HA-GPC6. ****p < 0.0001, Welch’s t-test, N = 3 animals, n = 49 HA-GFP, 47 HA-GPC6-expressing cells. **o-r**: Graphs quantifying neurons expression HA-GFP-MYC or HA-GPC6 comparing (**o**) Nup98 C/Nuclear envelope (NE) ratio (****p < 0.0001, Welch’s t-test, N = 3 animals, n = 78 HA-GFP-MYC, 77 HA-GPC6-expressing cells.), (**p**) Ran N/C ratio (****p < 0.0001,Welch’s t-test, N = 3 animals, n = 79 HA-GFP-MYC, 76 HA-GPC6-expressing cells.), (**q**) TDP-43 N/C ratio (****p < 0.0001, Welch’s t-test, N = 3 animals, n = 67 HA-GFP-MYC, 62 HA-GPC6-expressing cells.), (**r**) TDP-43 nuclear intensity (****p < 0.0001, Welch’s t-test, N = 3 animals, n = 67 HA-GFP-MYC, 62 HA-GPC6-expressing cells.). A.U.: arbitrary unit. All graphs: mean ± sem.

Ablation of GDE2 results in sustained canonical Wnt signaling in cortical neurons that disrupts NPC, NCT, and TDP-43 localization^29^. Specifically, Nucleoporin (Nup) 98, a structural component of the NPC that is required for maintaining the nuclear barrier and selective protein import and export, abnormally accumulates in the cytoplasm^51^. Moreover, the Ras-related GTPase (Ran) that controls the nuclear import and export of proteins through a gradient of nuclear-enriched Ran-GTP compared with cytoplasmic Ran-GDP is disrupted, leading to a reduction in Ran nuclear/cytoplasmic (N/C) ratio^52^. Nuclear import of TDP-43 utilizes the Ran gradient^12,53^, and consistent with disrupted NCT, TDP-43 N/C ratios and TDP-43 nuclear intensity are reduced upon GDE2 ablation and ectopic Wnt activation^29^. Strikingly, HA-GPC6-expressing neurons recapitulated these changes, in that they showed an increase in Nup98 cytoplasmic accumulation (Fig. 3c-f, o, Fig. S3a) and a reduction in the Ran N/C ratio (Fig. 3g-j, p, Fig. S3b) compared to neurons expressing HA-GFP-MYC. Further, neurons expressing HA-GPC6 showed a decrease in TDP-43 nuclear intensities and N/C ratio relative to neurons expressing control virus (Fig. 3k-n, q-r, Fig. S3c-d). Therefore, GPC6 overexpression in adult neurons phenocopies *Gde2*KO and ectopic Wnt activation by its ability to increase canonical Wnt signaling, disrupt NPC and NCT protein distribution, and promote TDP-43 cytoplasmic mislocalization and nuclear reduction.

### GPC6 genetic reduction rescues TDP-43 mislocalization in *Gde2*KO animals

Our previous observations demonstrate that increased GPC6 surface expression in cortical neurons is sufficient to activate canonical Wnt signaling and trigger NPC/NCT disruptions and TDP-43 mislocalization. We reasoned that if GDE2 is a physiological regulator of GPC6 surface expression that is important for this pathway, then reducing the levels of endogenous GPC6 should rescue these changes in *Gde2*KO animals. To test this hypothesis, we genetically reduced GPC6 in *Gde2*KO and *Gde2*KO;*Wnt-MYC-GFP* mice by generating *Gde2*KO;*Gpc6*+/-(*Gde2*KO;*Gpc6*Het) and *Gde2*KO;*Gpc6*Het;*Wnt-MYC-GFP* animals. We did not completely ablate GPC6 in these animals because *Gpc6*KO mice die immediately after birth^54^. Animals were aged to 4 months before analysis by Western blot and immunohistochemistry. Western blots of cortical lysates from *Gde2*KO;*Gpc6*WT and *Gde2*KO;*Gpc6*Het animals showed that this genetic strategy reduced GPC6 protein amounts in *Gde2*KO animals by approximately 30% (Fig. 4a-b). To determine whether GPC6 reduction can rescue canonical Wnt activation in cortical neurons of *Gde2*KO animals, we examined neuronal MYC expression in 4-month-old WT;*WNT-MYC-GFP*, *Gde2K*O:*WNT-MYC-GFP*, and *Gde2*KO;*Gpc6*Het;*WNT-MYC-GFP* animals. Compared to WT;*WNT-MYC-GFP* animals, *Gde2K*O;*WNT-MYC-GFP* mice showed a marked increase in the proportion of nuclear MYC+ neurons, consistent with aberrant neuronal Wnt activation upon GDE2 ablation (Fig. 4c, c’, d). In contrast, genetic reduction of *Gpc6* in *Gde2*KO;*Gpc6*Het;*WNT-MYC-GFP* animals reduced the proportion of nuclear MYC+ neurons normally seen in *Gde2K*O:*WNT-MYC-GFP* mice to WT:*WNT-MYC-GFP* levels (Fig. 4c, c’, d). No changes were observed in non-neuronal MYC+ cells between the three genotypes (Fig. 4c, c’, e), which is consistent with previous studies showing that GDE2 loss does not perturb canonical Wnt signaling in non-neuronal cells. Thus, these observations are consistent with the model that GDE2 regulates the surface expression of GPC6 to regulate canonical Wnt signaling in neurons.

**Figure 4:**
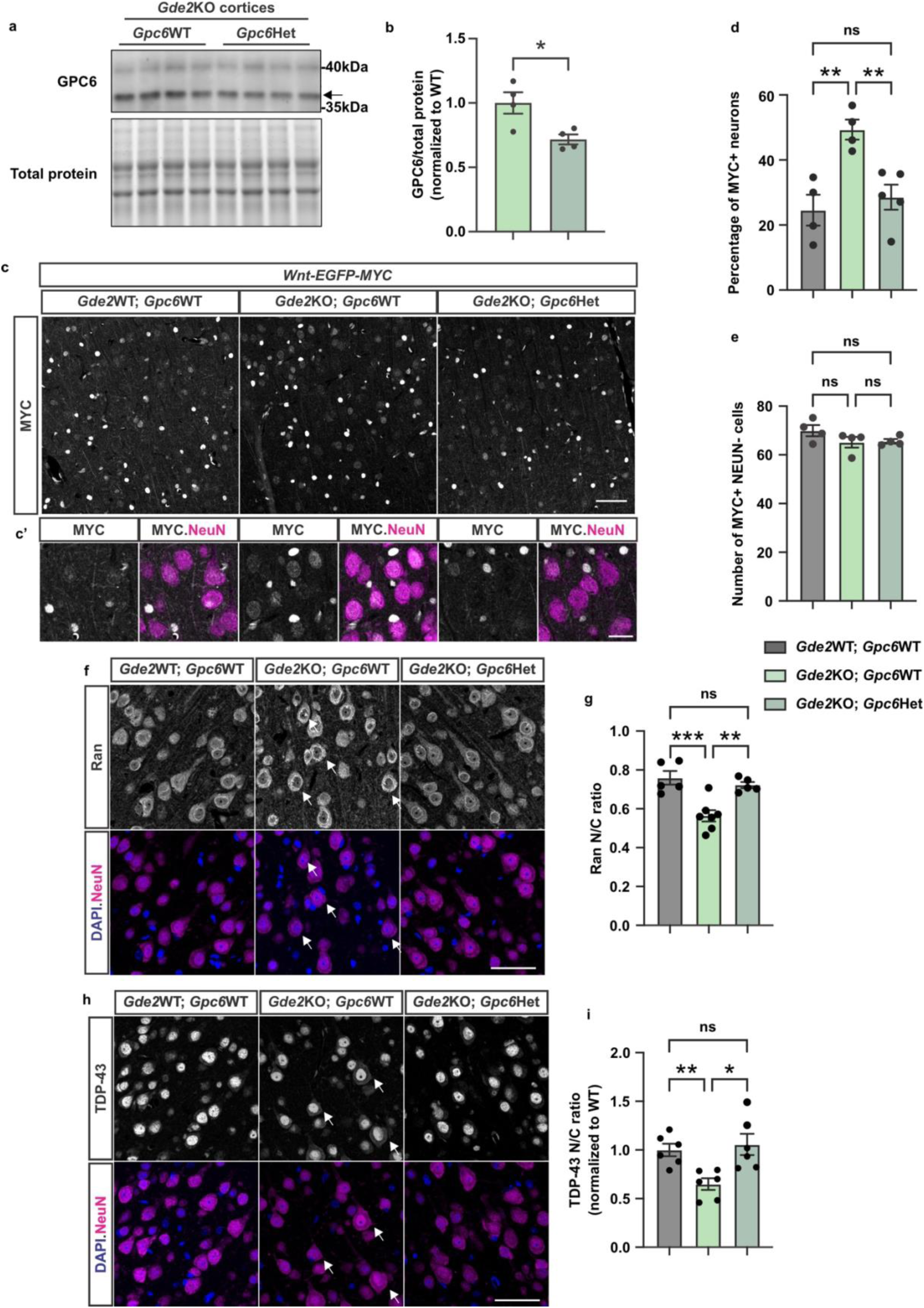
*Gpc6* genetic reduction restores NCT and TDP-43 nuclear localization in *Gde2*KO animals. **a-b.** Representative western blot images of GPC6 in *Gpc6*WT and *Gpc6*Het cortical extracts (**a**) and graph (**b**) quantifying GPC6 levels normalized to total protein. Arrow points to the band for furin cleaved GPC6. Gpc6WT vs. Gpc6Het: *p = 0.0215, n = 4, Unpaired t-test. **c-d**. Representative images of immunostained cortical sections of 4-month-old WT;*WNT-MYC-GFP*, *Gde2K*O:*WNT-MYC-GFP*, and *Gde2*KO;*Gpc6*Het;*WNT-MYC-GFP* animals (**c, c’**) and graphs quantifying the proportion of MYC+ neurons (**d**) and number of MYC+ non-neuronal cells per field of view (**e**) in the cortex. Images in **c’** are magnified from **c**. Scale bars: **c** = 50μm, **c’** = 20μm. **d**: WT;*WNT-MYC-GFP* vs. *Gde2K*O:*WNT-MYC-GFP*: **p = 0.0044; *Gde2K*O:*WNT-MYC-GFP* vs. *Gde2*KO;*Gpc6*Het;*WNT-MYC-GFP*: **p = 0.0097; WT;*WNT-MYC-GFP* vs. *Gde2*KO;*Gpc6*Het;*WNT-MYC-GFP*: ns p = 0.7573. N = 4-5 animals, n ∼ 600 neurons per animal. Ordinary one-way ANOVA with Tukey’s multiple comparisons test. e: WT;*WNT-MYC-GFP* vs. *Gde2K*O:*WNT-MYC-GFP*: ns p = 0.2294; *Gde2K*O:*WNT-MYC-GFP* vs. *Gde2*KO;*Gpc6*Het;*WNT-MYC-GFP*: ns p = 0.9888; WT;*WNT-MYC-GFP* vs. *Gde2*KO;*Gpc6*Het;*WNT-MYC-GFP*: ns p = 0.2794. N = 4-5 animals. **f, h:** Representative images of immunostained cortical sections of 4-month-old WT, *Gde2K*O, and *Gde2*KO;*Gpc6*Het. Scale bar = 50μm. Arrows highlight cells in *Gde2K*O exhibiting nuclear exclusion of Ran (**f**) and cytoplasmic mislocalization of TDP-43 (**h**). **g, i**. Graph quantifying Ran N/C ratios (**g**. WT vs. *Gde2*KO: ***p = 0.0007; *Gde2K*O vs. *Gde2*KO;*Gpc6*Het: **p = 0.0039; WT vs. *Gde2*KO;*Gpc6*Het: ns p = 0.6670. Ordinary one-way ANOVA with Tukey’s multiple comparisons test) and TDP-43 N/C ratios (**i**. WT vs. *Gde2*KO: **p = 0.0070; *Gde2K*O vs. *Gde2*KO;*Gpc6*Het: *p = 0.0314; WT vs. *Gde2*KO;*Gpc6*Het: ns p = 0.9577. Brown-Forsythe and Welch ANOVA tests). N = 6 animals, ≥120 cells per animal. All graphs: mean ± sem.

To determine if the dampening of Wnt activation by the genetic reduction of *Gpc6* is sufficient to rescue NCT and TDP-43 abnormalities in *Gde2*KO animals, we examined Ran and TDP-43 distribution in cortical neurons of 4-month-old WT, *Gde2K*O, and *Gde2*KO;*Gpc6*Het animals by immunohistochemistry. *Gde2*KO cortical neurons exhibit a reduction in the N/C ratio of Ran, consistent with the disruption of the Ran gradient important for NCT (Fig. 4f-g). Notably, *Gde2*KO;*Gpc6*Het mice showed an equivalent Ran N/C ratio to WT animals, suggesting that NCT trafficking is restored upon GPC6 reduction (Fig. 4f-g). In line with this observation, the reduction in TDP-43 N/C ratio observed in *Gde2*KOs, indicative of impaired TDP-43 localization, is rescued to WT levels in *Gde2*KO;*Gpc6*Het mice (Fig. 4h-i). In summary, these observations provide evidence that GDE2 regulates GPC6 expression to modulate neuronal Wnt signaling, which ensures the integrity of NCT and TDP-43 nuclear localization.

## Discussion

Previous studies have determined that GDE2 encodes a physiological pathway that prevents sustained canonical Wnt signaling in neurons to ensure NPC integrity, appropriate NCT, and nuclear TDP-43 expression and function^29^. We show here that GDE2 inhibits persistent canonical Wnt signal transduction in neurons by regulating the surface expression of GPC6. Excessive GPC6 expression *in vivo* potentiates Wnt activation, resulting in the disruption of the NPC, alterations in Ran-dependent NCT, and TDP-43 mislocalization, while genetic reduction of GPC6 rescues these cellular changes in *Gde2*KO animals. These observations highlight GPC6 as an important activator of canonical Wnt signaling in neurons, whose regulation by GDE2 is critical for preserving the NPC, NCT, and appropriate localization of TDP-43. These observations provide insight into fundamental, physiological pathways that are necessary for postmitotic neuronal function and viability. GDE2 dysfunction, GDE2-dependent Wnt activation, and GPC6 have been independently implicated in neurodegenerative diseases with TDP-43 abnormalities^26–29,37^. Our observations suggest the compelling possibility that GDE2, GPC6, and canonical Wnt pathway regulation integrate to form a physiologically important signaling axis that, when disrupted, contributes to NPC, NCT, and TDP-43 pathologies in disease.

Wnt signaling in mammals is highly complex, with 19 identified Wnt ligands, 2 LRP receptors, and 10 Fzd co-receptors^39^. We utilized a scRNA-seq dataset of the adult mouse cortex^38^ to identify Wnt components that mediate the regulation of Wnt signaling by GPC6, and focused on the brain-specific Wnt ligand, Wnt7b, Wnt receptor LRP6, and a subset of Fzd co-receptors that are co-expressed with GDE2 and GPC6 in the adult cortex. Using a cell line that reports canonical Wnt activity, we found that GPC6 potentiates Wnt activation elicited by Wnt7b when co-expressed with Fzd1, Fzd4, and Fzd7, but not Fzd8. GPC6 belongs to the Dlp-class of Glypicans^30^. Mechanistically, Dlp-class Glypicans shield the lipid moiety of the Wnt ligand and facilitate their transfer to Fzd receptors, thus acting as a reservoir of signaling-competent Wnt ligand at the cell surface^31,32^. Our study suggests that there is some specificity in the mechanism of GPC6-dependent transfer of Wnt ligands to Fzd co-receptors. However, the basis of this specificity is unclear. Future studies that interrogate the structural interactions between GPC6, Wnt ligands, and Fzds may clarify this issue. Our analysis of GDE2 and GPC6 regulation of Wnt signaling focuses primarily on deep-layer cortical neurons. Notably, GPC1 and GPC4 are also co-expressed with GDE2 in deep-layer neurons, albeit at much lower levels than GPC6. It will be interesting to examine the contributions of GPC1 and GPC4 to GDE2-dependent Wnt activation and determine whether these pathways, in addition to GPC6, extend to other neuronal subtypes in the brain. Further molecular insight can be gleaned by determining the *in vivo* contributions of Fzd1, Fzd4, and Fzd7 to GDE2/Glypican-mediated Wnt7b signal transduction in different cellular contexts.

Studies of *WNT-MYC-GFP* mice here and elsewhere suggest that cortical neurons exhibit low and transient activation of canonical Wnt signaling *in vivo*^39,55,56^. A key feature of Dlp-class Glypicans that facilitates ligand transfer to Wnt receptors at the cell surface is the GPI-anchor, which enables their rapid diffusion through the plane of the membrane^30^. We propose that GDE2 GPI-anchor cleavage of GPC6 provides an effective and rapid means to slow/prevent ligand transfer to ensure low levels of Wnt signaling in cortical neurons. This is critical to prevent a buildup of GPC6 surface expression that would increase and sustain Wnt activation, which would negatively impact neuronal health and viability due to downstream effects on the NPC, NCT, and TDP-43 nuclear localization and function. Importantly, complete blockade of Wnt signaling is detrimental, as transient activation of Wnt signaling in neurons is required for neuronal functions such as dendritic plasticity^55,56^. Accordingly, we propose that periodic downregulation of GDE2 activity is essential to preserve the dynamics of neuronal Wnt activation. Studies of GDE2 developmental functions reveal that thiol-redox mechanisms regulate GDE2 trafficking to the cell surface and surface activity^57,58^. This suggests the compelling possibility that similar pathways might act physiologically to regulate GDE2 surface activity in adult neurons; however, this requires further investigation.

Studies in animal models of disease, human postmortem brain, cerebrospinal fluid, and iPSC-derived neurons (iSNs) suggest that GDE2 dysfunction and abnormal activation of Wnt signaling contribute to TDP-43 mislocalization and impaired nuclear function in ALS and ALS/FTD^28,29^. Our discovery that GDE2 regulation of GPC6 is a physiological pathway that prevents sustained Wnt activation in neurons suggests potential contributions of GPC6 to diseases with TDP-43 proteinopathies in the context of GDE2 dysfunction/Wnt activation. Interestingly, GPC6 is associated with AD and ADRDs in GWAS and is found to be mislocalized in spinal cord tissue of patients with ALS^33–37^. Further, GPC6 release is reduced in the *SOD1G93A* model of ALS^26^, and work in *Drosophila* suggests *Dlp* is a target of TDP-43 proteinopathy, with reduced expression at the neuromuscular junction (NMJ)^37^. These observations, combined with our study, suggest that GPC6 is likely to have complex and varied contributions to neurodegenerative disease onset, progression, and pathology.

## Methods

### Animals

Mice were bred and maintained in accordance with approved Johns Hopkins University Institutional Animal Care and Use Committee (JHU IACUC) protocols. *Gde2*KO^24^, WT;*Wnt-MYC-GFP* (*Rosa26 Tcf/Lef H2B-EGFP-6xMYC*)^45^, and *Gpc6+/-*^54^ mice were bred, maintained, and genotyped as described previously. All experimental protocols were approved by JHU IACUC. All experiments were carried out in accordance with relevant guidelines and regulations, including JHU IACUC and the Animal Research: Reporting of In Vivo Experiments (ARRIVE) guidelines. For each experiment, the groups being compared, the respective experimental units, and the sample sizes (number of cells and animals analyzed) are listed in the figure legends. No statistical power analysis was used to predetermine sample size. Sample sizes were determined to be similar or to exceed those previously reported in the literature. There were no exclusions of animals in this study. No randomization sequence was generated or used to allocate animals to control or treatment groups. However, animals were randomly picked for injections, and AAV injections were performed alternating between control and AAV GPC6 to minimize confounding effects from the order of treatments. Animals with different treatments or genotypes were maintained in mixed housing, separated by sex, until the age/timepoint listed in the figure legends for each experiment. Both males and females were used in all experiments, and the genotypes for each experiment are listed in the figures and figure legends. The statistical methods used are included in the figure legends. Datasets were tested for assumptions in their distribution, and corrections were made in statistical testing if the assumptions were not met.

### AAV vector construction and injections

All AAV plasmid backbones were based on AAV-GFP.Cre^59^ (Addgene, 49056). AAV-GFP was constructed by deleting the coding sequence of Cre. For AAV-HA.GFP.MYC and AAV-HA.GPC6, the coding sequences were synthesized from pRK-HA.GFP.MYC (Addgene, 137763) and pDisplay HA.GPC6^16^, respectively, and subcloned into the AAV backbone by replacing GFP/Cre. AAV-HA.GFP was constructed by deleting the MYC tag. The AAV vectors were packaged by Janelia Viral Tools using the PHP.eB capsid^60,61^.

AAV vectors were delivered to mice retro-orbitally at p28 at 10^11^ viral genome per animal as previously described^60^. Prior to retroorbital injections, animals weighed around 15 grams and were anesthetized by intraperitoneal injection of 0.01ml/g Avertin (1.3% 2,2,2-Tribromoethanol (T48402-25G) and 0.7% 2-methyl-2-butanol (Sigma 240486) in Phosphate Buffered Saline (PBS)). Animals were monitored post-injection. Prior to tissue harvesting at the time points indicated, animals were euthanized by intraperitoneal injection of 0.02ml/g Avertin (see also Immunohistochemistry).

### Immunohistochemistry (IHC)

4-month-old animals weighed around 25 grams prior to tissue harvesting, and there were no gross weight differences between genotypes. Mice were anesthetized with 0.02ml/g Avertin solution (1.3% 2,2,2-Tribromoethanol (T48402-25G) and 0.7% 2-methyl-2-butanol (Sigma 240486) in Phosphate Buffered Saline (PBS)) by intraperitoneal injection before transcardial perfusion with 0.1M Phosphate Buffer (PB) and 4% Paraformaldehyde (PFA) in 0.1M PB. Brains were dissected, post-fixed in 4% PFA for 18-20 hours, washed with PBS, and prepared for embedding in cryomolds or paraffin blocks as previously described^26^.

Cryo-embedded brains were sectioned on a cryostat (Leica CM3050S) at 30-40µm. IHC-immunofluorescence (IF) was performed on free-floating sections. Paraffin blocks were cut into 4µm sections on a Rotary microtome (Leica RM2235) and collected on slides. Paraffin sections were deparaffinized with Xylenes and rehydrated in an ethanol series immediately prior to staining. Immunostaining was performed as previously described^29^. Briefly, sections were washed in PBS and permeabilized in 0.3% Triton-X-100 in PBS (PBST). For paraffin sections, antigen retrieval was performed with 0.1M sodium citrate buffer (pH 6) for 20 minutes in a 95°C water bath. Sections were then incubated for at least 1 hour with blocking solution (5% Normal Donkey Serum (NDS) or 5% Bovine Serum Albumin (BSA) (Sigma, A9647-100G) in PBS and 0.3% Triton-X-100 in PBS). Sections were incubated overnight with primary antibodies at 4°C and washed with PBS the next day. Sections were then incubated with the appropriate fluorescently conjugated secondary antibodies (Jackson Immunoresearch) for 1-2 hours at room temperature (RT). Nuclei were stained with Hoechst 33342 (1:500, Thermo Fisher) in PBS for 15 minutes at RT. Sections were mounted on slides with Prolong Gold mounting media (ThermoFisher, P36931) and coverslipped before imaging. Images were acquired with a Zeiss LSM 700 microscope. The same settings were used for all images acquired within the same experiment. See Table 1 for antibody information.

**Table 1.** Antibody Information.

| Primary Antibodies |  |  |  |
| --- | --- | --- | --- |
| Antibody | Source | Catalog number | Application and concentration |
| GFP | Invitrogen | A11120 or A11122 | IHC: 1:500 |
| GPC6 | R&D systems | AF2845 | Immunoblotting: 1:2500 |
| Mouse GDE2 | Covance |  | Immunoblotting: 1:1000 |
| HA (6EA) | Cell Signaling Technologies | 2367S | IHC: 1:500<br>Immunoblotting: 1:2500 |
| HA (C29F4) | Cell Signaling Technologies | 3724 | IHC: 1:500 |
| hFAB Rhodamine anti-Actin | BioRad | 12004163 | Immunoblotting: 1:2500 |
| hFAB Rhodamine anti-Tubulin | BioRad | 12004166 | Immunoblotting: 1:5000 |
| MYC | Cell Signaling Technologies | 2278 | IHC: 1:250 |
| NeuN | Synaptic Systems | 266 004 | IHC: 1:1000 |
| non-phospho-β-catenin | Cell Signaling Technologies | 8814 | Immunoblotting: 1:1000 |
| Nup98 | Abcam | ab50610 | IHC: 1:250 |
| Ran | BD Transduction | 610341 | IHC: 1:500 |
| TDP-43 | Proteintech | 10782-2-AP | Immunoblotting: 1:5000<br>IHC: 1:500 |

### RNAscope

RNAscope coupled with IHC was carried out using the RNAscope Multiplex Fluorescent v2 Assay with the Co-Detection kit (ACDbio, 323110) according to the manufacturer’s protocol. *Gpc6* mRNA was virtualized using the probe Mm-Gpc6-C2 (ACDBio, Cat No. 442841-C2) and Opal-570 fluorophore (Akoya Biosciences, FP1488001KT, 1:500), and costained with NeuN (synaptic systems, 266004,1:500). Slides were mounted with ProLong Gold mounting media with DAPI (ThermoFisher, P36931), coverslipped, and imaged on a Zeiss LSM800 confocal microscope.

### Quantitative Image Analysis

All images were quantified using ImageJ software (NIH) as previously descirbed^29^. For mouse tissues, 2-4 sections per mouse were analyzed depending on the experiment. For fluorescent stains, threshold-based cell counting macros created in ImageJ were used to quantify NeuN-positive cells and cells double positive for both NeuN and MYC. Neurons were considered positive for MYC if their nuclear intensity was more than 2x standard deviations above the mean background intensity.

To quantify nuclear/cytoplasmic (N/C) or C/nuclear envelope (NE) ratios, DAPI was used to delineate the nucleus from the cytoplasm in neurons. NeuN signal outside of DAPI was used to define the cytoplasm. The nuclear and cytoplasmic intensities were measured as the average fluorescence intensities from three regions of interest (ROIs) of the same size in the nucleus and in the cytoplasm in the channel of interest. The NE intensity was measured by drawing a freehand line of 2-pixel width around the nucleus. Five background measurements were taken per image, the average of which was used to correct for differences in background intensity. The final N/C ratio was calculated by dividing the average background-corrected nuclear intensity by the average background-corrected cytoplasmic intensity for each cell. The final C/NE ratio was calculated by dividing the average background-corrected cytoplasmic intensity by the average background-corrected NE intensity for each cell. Nuclear intensities were calculated using a mask of the area within DAPI to measure mean intensity in the relevant channel of interest.

### Cell culture

#### Mouse primary cortical neuronal culture

P0 or P1 pups were euthanized by cryoanesthesia followed by decapitation. Mouse primary cortical cultures were prepared from the extracted mouse cortices and plated on poly-l-lysine-coated plates or acid-washed coverslips as previously described^27^ with minor modifications. Cells were maintained in Neurobasal medium supplemented with 2% B27, 1% L-glutamine, and 1% Pen/Strep. 5µM cytosine arabinoside was added on DIV2 to inhibit glial growth and removed on DIV3. From DIV4, cultures were fed every 3 days and maintained at 37°C until harvest at DIV21.

#### Immortalized cell lines

SH-SY5Y (CRL-2266) cells were obtained from American Type Culture Collection (ATCC) and cultured according to the manufacturer’s protocol. SuperTopflash (STF) cells were a gift from Dr. Jeremy Nathans (Johns Hopkins University). Plasmids were transfected with FuGENE HD (Promega E2311) for STF cells and Lipofectamine 2000 (Invitrogen 11668-027) for SH-SY5Y cells accordingly to the manufacturer’s protocol. The plasmid expressing HA-GPC6 was previously described^16^. Plasmids expressing Wnt7b, Lrp6, Fzd1, Fzd3, Fzd4, Fzd7, and Fzd8 are kind gifts from Dr. Jeremy Nathans. siRNAs were purchased from Santa Cruz Biotechnology to knock down *hGPC6* (sc-75152) compared to non-targeting control (sc-37007).

### Cell Surface Biotinylation and Fractionation

Cell surface biotinylation and fractionation were performed according to a previously described protocol^62^ with minor modifications. Briefly, culture plates were placed on ice for 15 minutes prior to washing to prevent cold-shock-induced lysis. Cells were rinsed once with ice-cold PBSCM (PBS-calcium-magnesium: 1× PBS, 1mM MgCl^2^, 0.1mM CaCl^2^ (pH 8.0)), and then incubated with Sulfo-NHS-SS-biotin (1.0 mg/mL, Thermo Scientific 21331) for 30 min at 4°C. Cells were then washed with PBSCM and incubated in 20 mM glycine in PBSCM twice for 5 min to quench unreacted biotinylation reagent. Neurons were lysed in radioimmunoprecipitation assay (RIPA) buffer containing 1x protease inhibitor cocktail (Sigma, P8340) and spun down at 21,000g for 15 minutes at 4°C. Protein levels were measured using a BCA Protein Assay kit (ThermoFisher, 23225). Equal amounts of proteins were incubated overnight with NeutrAvidin agarose beads (ThermoFisher, 29201) at 4°C on a rotator. The beads were then washed four times with RIPA buffer containing protease inhibitor, and the captured biotinylated fraction was eluted by incubating the beads in 2x Laemmli buffer at 95°C for 10 minutes. The total and surface fractions were then analyzed using immunoblotting.

### Immunoblotting

4-month-old animals weighed around 25 grams prior to tissue harvesting, and there were no gross differences between genotypes. Mice were anesthetized with 0.02ml/g Avertin solution (1.3% 2,2,2-Tribromoethanol (T48402-25G) and 0.7% 2-methyl-2-butanol (Sigma 240486) in Phosphate Buffered Saline (PBS)) by intraperitoneal injection before decapitation and tissue harvesting. Mouse cortical tissues were sonicated in RIPA buffer containing 1x protease inhibitor cocktail (Sigma, P8340) and spun down at 21,000g for 20 minutes at 4°C. Protein levels were standardized across all samples using a BCA Protein Assay kit (ThermoFisher, 23,225) and 4x Laemmli buffer was added to samples to a final concentration of 1x. Cultured cells were lysed directly in 1x Laemmli buffer, sonicated, and spun down at 21,000g for 10 minutes at RT. Immunoblotting was performed as previously described^29^. Samples were boiled and run on 7.5% or 10% polyacrylamide gels in tris/glycine buffer before transferring to polyvinylidene difluoride (PVDF) membranes. PVDF membranes were blocked with 5% milk in tris-buffered saline containing 0.3% Tween-20 (TBST) or Everyblot blocking buffer (Bio-Rad laboratories,12010020) for 1-2 hours at RT before applying primary antibodies overnight at 4°C. After washing with TBST, the membranes were incubated with the appropriate horseradish peroxidase (HRP) or fluorescent protein-conjugated secondary antibodies for 1 hour at RT. Membranes were washed again with TBST, developed using enhanced chemiluminescence substrate when appropriate (Kindle Biosciences, R1004), and imaged using a ChemiDoc Imager (Bio-Rad). Blots were analyzed using ImageJ software (NIH). See Table 1 for antibody information.

### TOPFlash Luciferase Assay

Renilla luciferase control construct (pRL-TK) was purchased from Promega (E2241). Dual-luciferase reporter assay was performed according to the manufacturer’s instructions (Promega, E1910). The readings of Firefly and Renilla luminescence were recorded using a Turner BioSystems Luminometer (TD-20/20). Relative luciferase activity was calculated by normalizing Firefly/Renilla values.

### Analysis of publicly available scRNAseq datasets

scRNAseq data of the adult mouse cortex and hippocampus from the Allen Brain Atlas^38^ were retrieved using CellxGene Census^63^ using the accession ID d7291f04-fbbb-4d65-990a-f01fa44e915b and visualized using the Scanpy^64^ and anndata^65^ packages in Python. No custom code was used in preparation of this manuscript.

### Statistical Analyses

Data were analyzed and plotted using GraphPad Prism (version 10). Statistical significance for pairwise comparisons was derived using a two-tailed Student’s t test with relevant corrections. For multiple comparisons, we used ANOVA with corrections for multiple comparisons. All values are reported as mean ± s.e.m. In figures, asterisks denote statistical significance: *P < 0.05, **P < 0.01, ***P < 0.001, ****P < 0.0001. Specific statistical information for each experiment is included in the figure legends.

## Data availability statement

The scRNAseq data sets analyzed in this study are available at https://portal.brain-map.org/atlases-and-data/rnaseq/mouse-whole-cortex-and-hippocampus-10x, and were retrieved using CellxGene Census^63^ using the accession ID d7291f04-fbbb-4d65-990a-f01fa44e915b. No other datasets were generated or analyzed during the current study.

## Acknowledgements

The authors thank Y. Li for technical assistance; Dr. J. Nathans for the *Wnt-MYC-GFP* mice, the SuperTopflash cell line, and expression vectors for Wnt ligands and Fzd receptors; Phillip Smallwood for help with Topflash luciferase assays; Dr. V. Gradinaru (CalTech) for AAV-PHP.eB rep-cap plasmids; Dr. L Goff for bioinformatics support. AAV-GFP/Cre was a gift from Dr. Fred Gage.

## Author contributions

N.Z. and S.S. conceived the study, designed experiments, interpreted data, and wrote the manuscript. N.Z. performed the experiments and formal analysis.

