## Supplementary Information for "Regulation of Glypican 6-mediated Wnt activation maintains TDP-43 nuclear localization in neurons"

Nan Zhang[1,2] and Shanthini Sockanathan[1*]

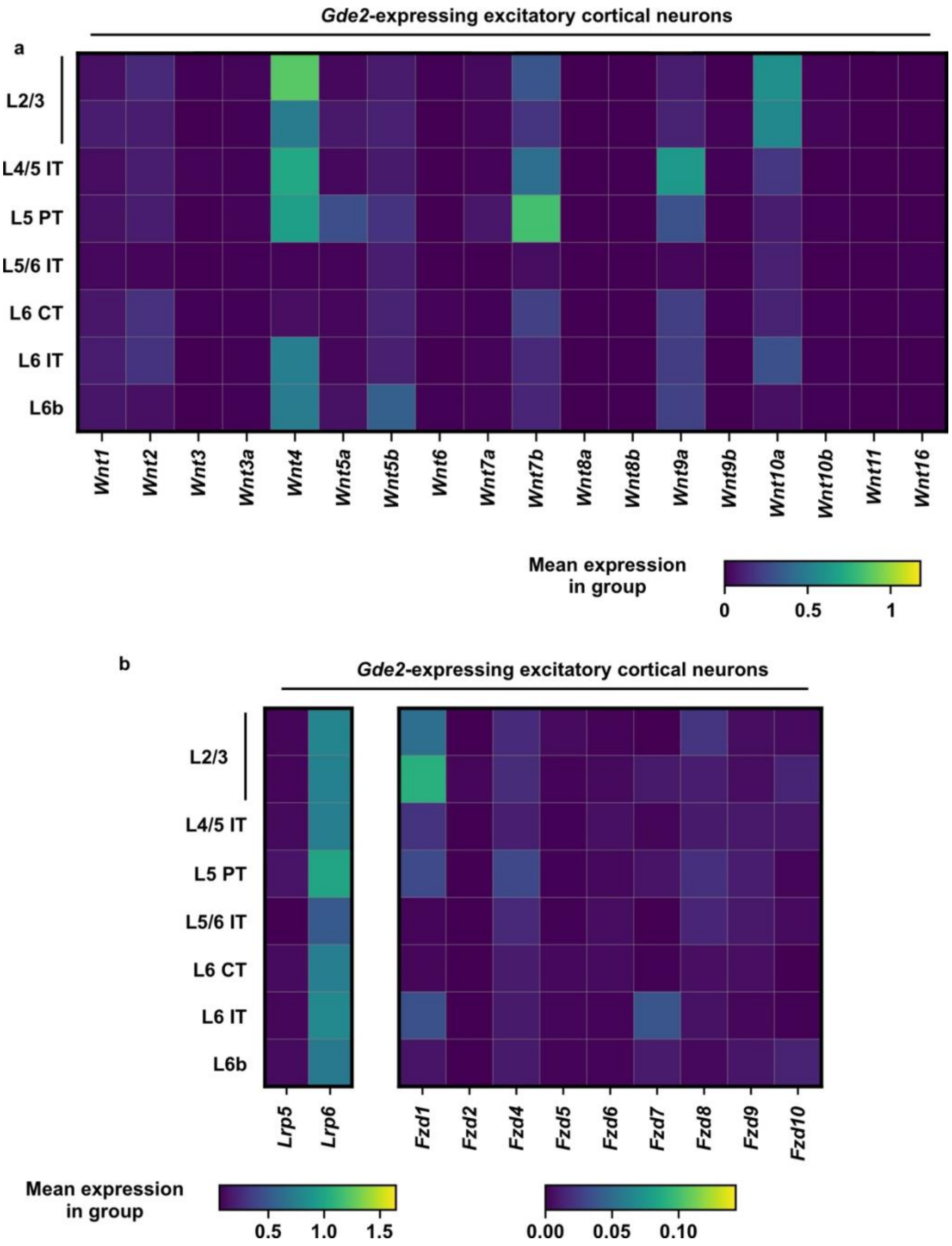


**Figure S1. The expression of Wnt signaling components in *Gde2*-expressing cortical neurons.**

**a-b**. Heat maps showing the expression of Wnt ligand-encoding genes, *Lrp5* and *Lrp6*, and *Fzd* family genes in *Gde2*-expressing neurons in the mouse cortex. Each subpanel is scaled differently to highlight the range of expression levels. Data from the Allen Brain Atlas. L: layer.

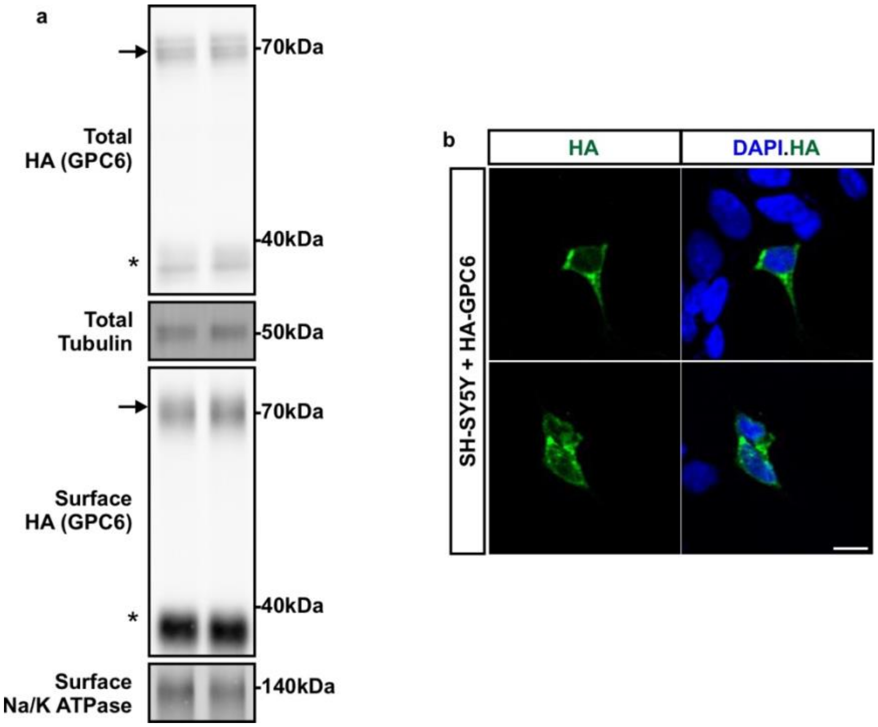


**Figure S2. Validation of HA-GPC6.**

**a**. Representative western blot images of surface biotinylation of SH-SY5Y cells overexpressing HA-GPC6 demonstrating that HA-GPC6 is expressed at the expected molecular weight and localized on the cell surface. The arrow marks the full-length form of HA-GPC6; asterisk marks HA-GPC6 after furin cleavage. **b**. Representative images of immunostained SH-SY5Y cells showing the localization of HA-GPC6. Scale bar = 10μm.

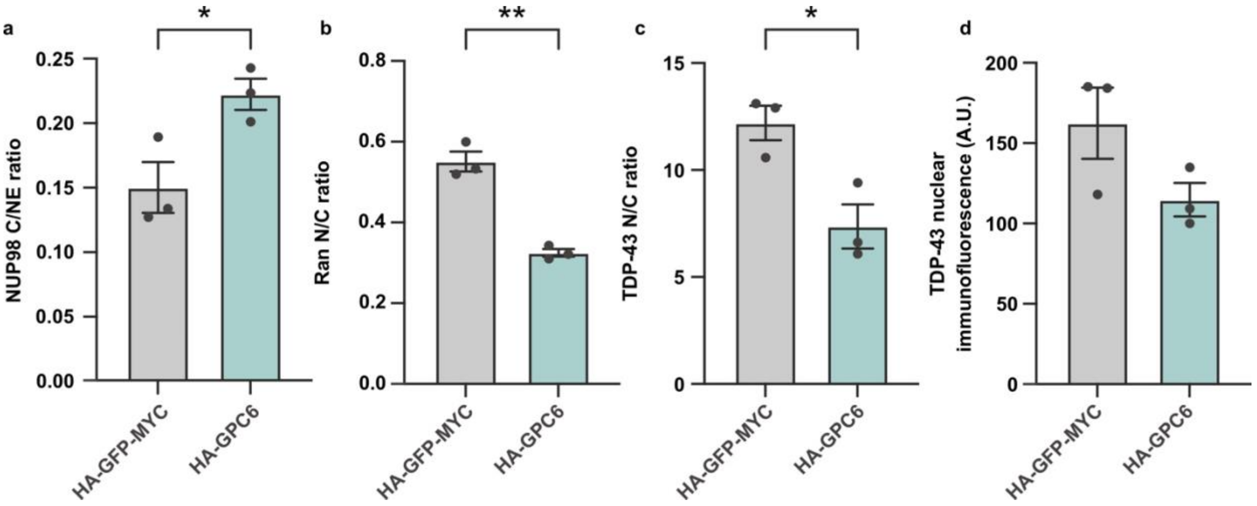


**Figure S3. GPC6 overexpression activates Wnt signaling and drives NPC, NCT, and TDP-43 abnormalities.**

Graphs quantifying neurons expressing HA-GFP-MYC or HA-GPC6 comparing by animal (**a**) Nup98 C/Nuclear envelope (NE) ratio (*p = 0.0202, paired t-test. N = 3 animals, n = 78 HA-GFP-MYC, 77 HA-GPC6-expressing cells.), (**b**) Ran N/C ratio (**p = 0.0057, paired t-test. N = 3 animals, n = 79 HA-GFP-MYC, 76 HA-GPC6-expressing cells.), (**c**) TDP-43 N/C ratio (*p = 0.0405, paired t-test. N = 3 animals, n = 67 HA-GFP-MYC, 62 HA-GPC6-expressing cells.), (**d**) TDP-43 nuclear intensity (ns p = 0.1041, paired t-test. N = 3 animals, n = 67 HA-GFP-MYC, 62 HA-GPC6-expressing cells.). A.U.: arbitrary unit. All graphs: mean ± sem.

### Uncropped immunoblot images

Figure 1d:

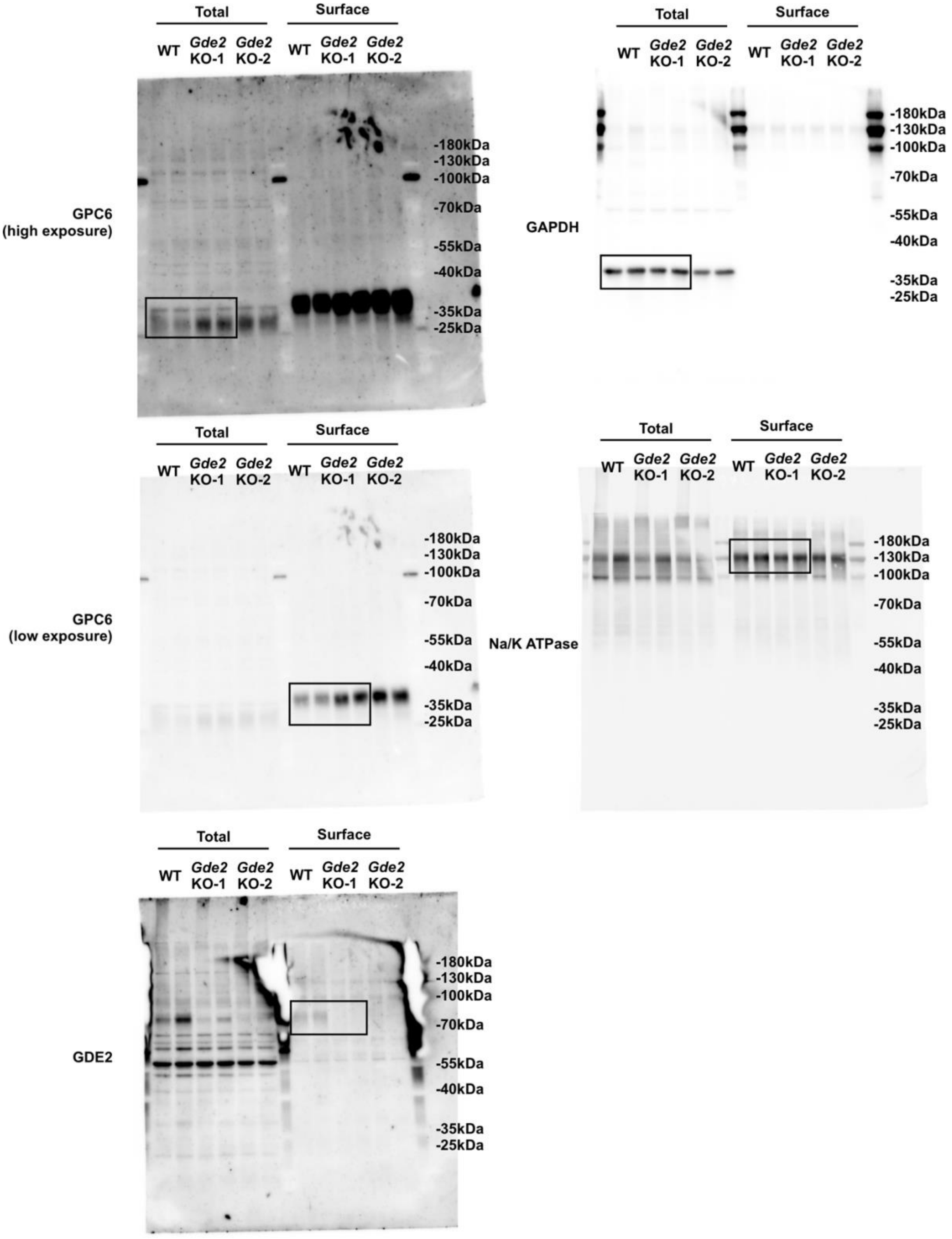

Figure 2a:

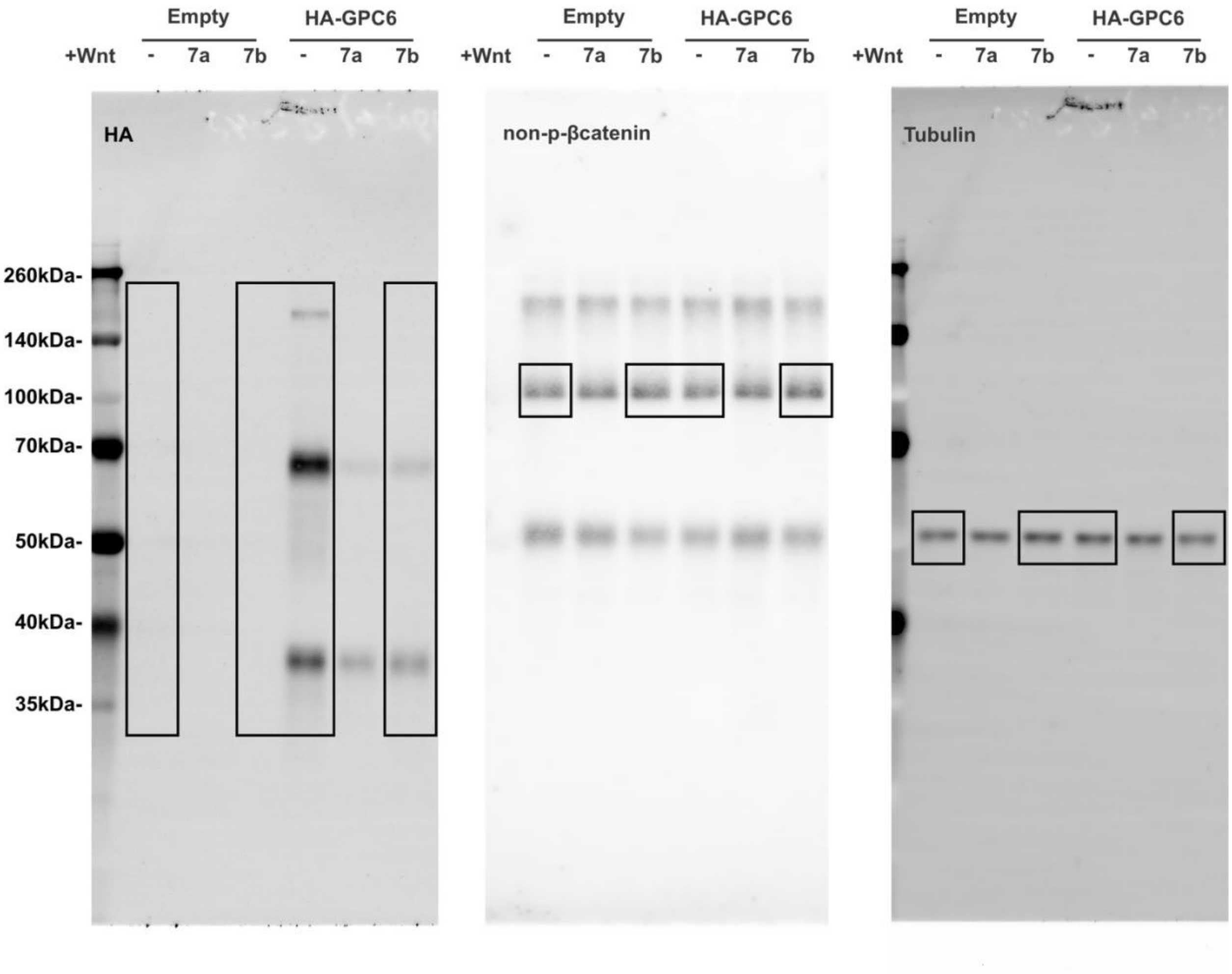
Empty
HA-GPC6
+Wnt
-
7a
7b
-
7a
7b
HA
260kDa-
140kDa-
100kDa-
70kDa-
50kDa-
40kDa-
35kDa-
Empty
HA-GPC6
+Wnt
-
7a
7b
-
7a
7b
non-p-βcatenin
Empty
HA-GPC6
+Wnt
-
7a
7b
-
7a
7b
Tubulin

Figure 2d:

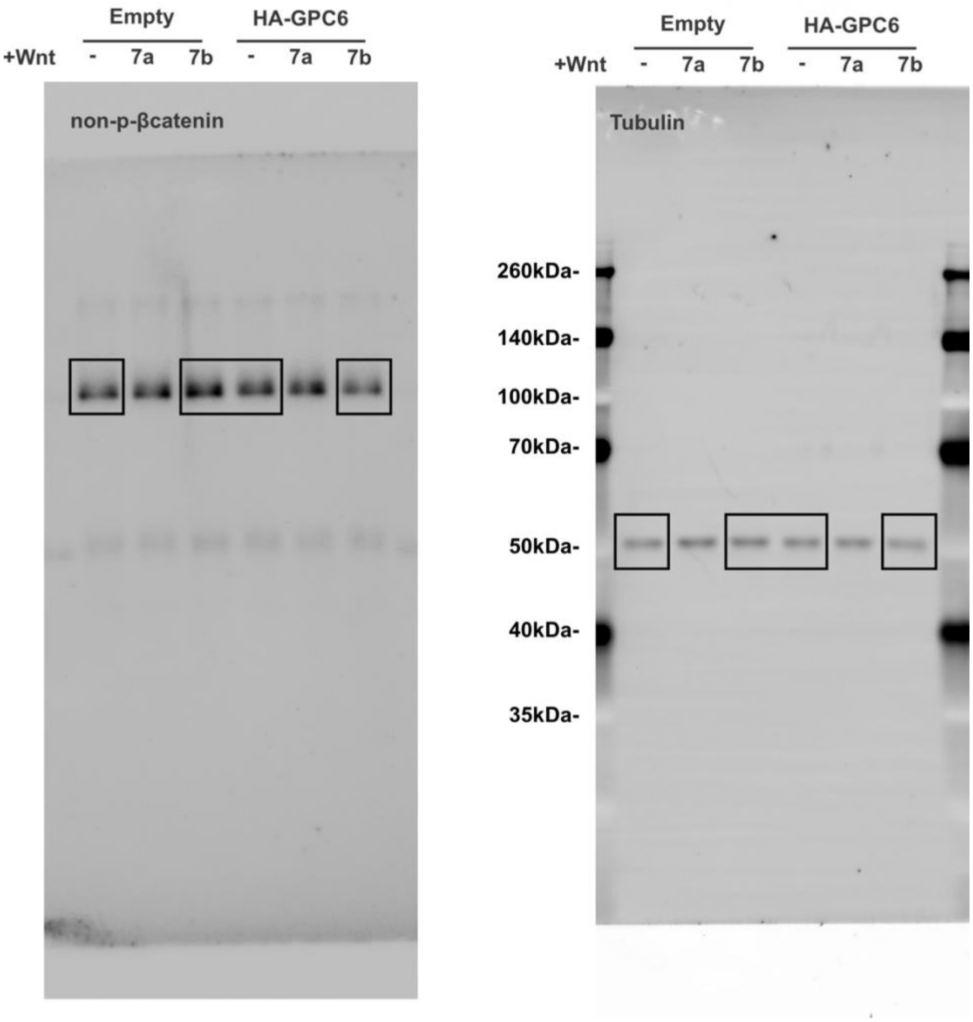
Empty
HA-GPC6
+Wnt
-
7a
7b
-
7a
7b
non-p-βcatenin
Empty
HA-GPC6
+Wnt
-
7a
7b
-
7a
7b
Tubulin
260kDa-
140kDa-
100kDa-
70kDa-
50kDa-
40kDa-
35kDa-

Figure 4a:

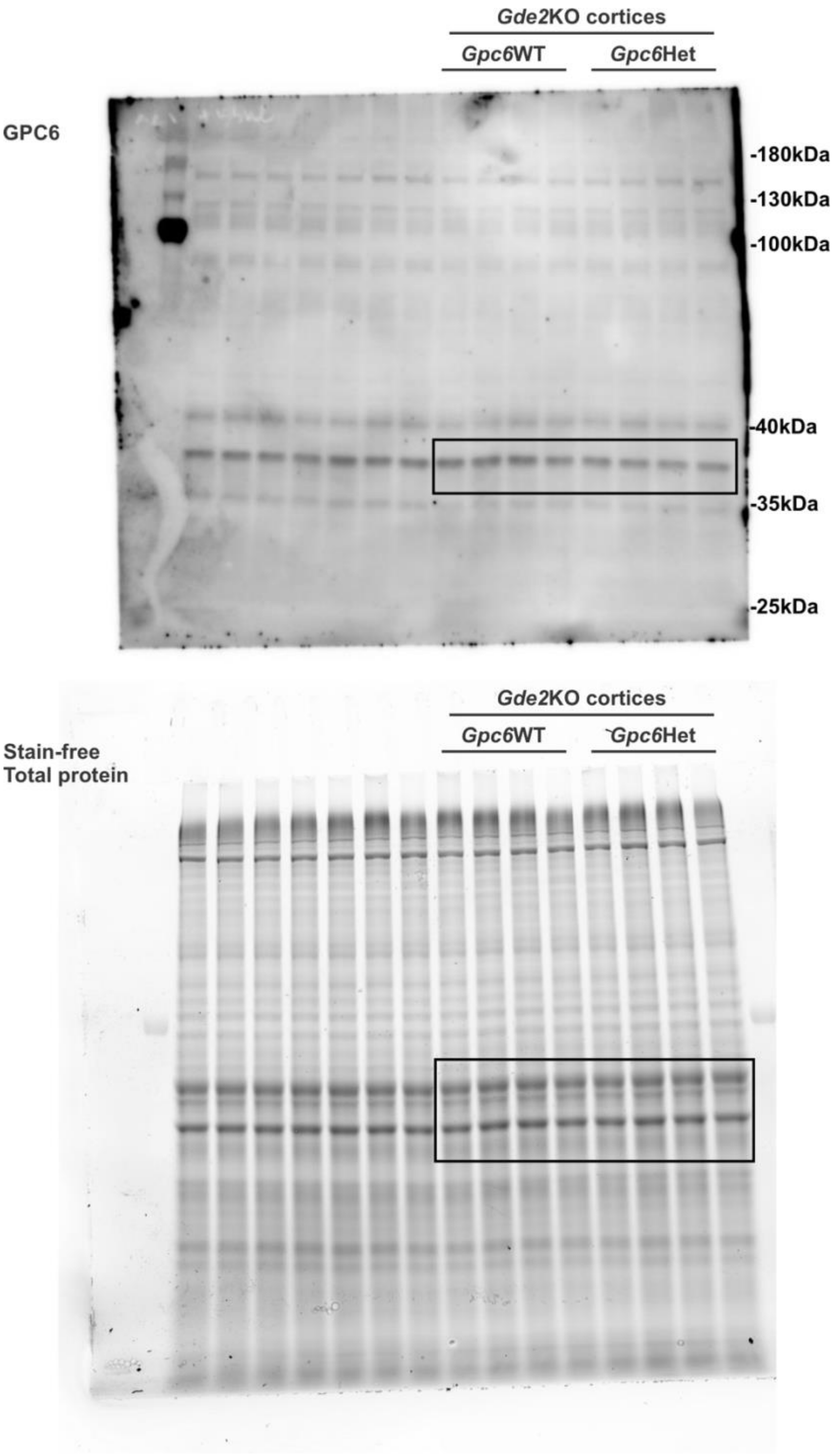

Gde2KO cortices
Gpc6WT
Gpc6Het
GPC6
-180kDa
-130kDa
-100kDa
-40kDa
-35kDa
-25kDa
Gde2KO cortices
Gpc6WT
Gpc6Het
Stain-free
Total protein
